# Kaposi sarcoma-associated herpesvirus infection in HIV patients: potential role of HIV-associated extracellular vesicles

**DOI:** 10.1101/640532

**Authors:** Lechuang Chen, Zhimin Feng, Guoxiang Yuan, Benjamin Reinthal, Fengchun Ye, Ge Jin

## Abstract

Kaposi sarcoma-associated herpesvirus (KSHV) is the causal agent for Kaposi sarcoma (KS), the most common malignancy in people living with HIV/AIDS. The oral cavity is a major route for KSHV infection and transmission. However, how KSHV breaches the oral epithelial barrier for spreading to the body is not clear. Here we show that extracellular vesicles (EVs) purified from saliva of HIV-positive individuals and secreted by HIV-1-infected T cells promote KSHV infectivity in both monolayer and 3-dimensional models of immortalized and primary human oral epithelial cells, establishing the latency of the virus. The HIV trans-activation response (TAR) element RNA in HIV-associated EVs contributes to the infectivity of KSHV through the epidermal growth factor receptor (EGFR). Cetuximab, a monoclonal neutralizing antibody to EGFR, blocks HIV-associated EV-enhanced KSHV infection. Our findings reveal that saliva containing HIV-associated EVs is a risk factor for enhancement of KSHV infection and that inhibition of EGFR serves as a novel strategy for controlling KSHV infection and transmission in the oral cavity.

**Author summary:** Kaposi sarcoma-associated herpesvirus (KSHV) is a causal agent for Kaposi sarcoma (KS), the most common malignancy in HIV/AIDS patients. Oral transmission through saliva is considered the most common route for spreading of the virus among HIV/AIDS patients. However, the role of HIV-specific components in co-transfection of KSHV is unclear. We demonstrate that extracellular vesicles (EV) purified from saliva of HIV patients and secreted by HIV-infected T cells promote KSHV infectivity in immortalized and primary oral epithelial cells. HIV-associated EVs promote KSHV infection depends on the HIV trans-activation element (TAR) RNA and EGFR of oral epithelial cells, both can be targeted for reducing KSHV infection. These results reveal that HIV-EVs is a risk factor for KSHV co-infection in the HIV-infected population.

## Introduction

Kaposi sarcoma (KS), the most common malignancy in patients infected with HIV, is etiologically associated with infection by Kaposi sarcoma-associated herpesvirus (KSHV), also known as human herpesvirus 8 (HHV-8) [1]. This oncogenic gamma-herpesvirus is also linked with primary effusion lymphoma (PEL), multicentric Castleman’s disease (MCD), and KSHV inflammatory cytokine syndrome (KICS) in aging people and immune compromised adults [2]. Oral transmission of KSHV through saliva in particular is believed to be the most common route for spreading of the virus among homosexual people and “mother to child” transmission [3,4].

The oral mucosa has shown to be the first target of KSHV infection once the virus is in the oral cavity [5,6,7,8,9]. Although KS incidence has dramatically decreased in developed countries in the era of antiretroviral therapy (ART), KS remains the most frequent tumor in the HIV-infected population worldwide [10,11,12]. The oral milieu of HIV-infected patients has long been deduced to favor KSHV infection; however, the role of HIV in KSHV infection and transmission is largely unknown.

Most types of cells can release lipid membrane-enclosed vesicles, generally called extracellular vesicles (EVs), into the extracellular space and body fluids. Saliva and other body fluids contain a variety of EVs [13,14,15,16]. EVs are highly heterogeneous and dynamic and can be generally grouped into exosomes [17,18,19,20], macrovesicles [21], and apoptotic bodies based on biogenesis and the origin of vesicles [22]. EVs contain molecular components of their cells of origin, including proteins and RNAs, to play roles in intercellular communication, molecular transfer, and immune regulation at local and distant sites [13,20,23]. EVs derived from culture supernatants of latently HIV-1-infected T-cell clones do not contain HIV-1 viral particles, although these EVs do contain viral proteins such as Gag and the precursor form of Env protein (p160) [24]. The HIV transactivation response (TAR) element RNA, a precursor of several HIV-encoded miRNAs, can fold in the nascent transcript and facilitate binding of the viral transcriptional trans-activator (Tat) protein to enhance transcription initiation and elongation of HIV [25]. EVs isolated from HIV-1-infected cells or from HIV-positive patient sera contain TAR RNA in vast excess of total viral RNA [24,26]. TAR RNA-bearing EVs can induce proinflammatory cytokines in primary macrophages [27] and stimulate proliferation, migration and invasion of head and neck and lung cancer cells in an epidermal growth factor receptor (EGFR)-dependent manner [26]. EVs in body fluids of HIV/AIDS patients may mediate HIV-1 RNA and protein trafficking and affect HIV pathogenesis [28,29]. However, the role of salivary HIV-associated EVs in co-infection of KSHV has not been explored [29].

Here, we report that EVs purified from the saliva of HIV-infected patients and secreted from culture medium of latently HIV-infected T cells enhance KSHV infectivity in human oral epithelial cells cultured in both monolayer and 3-dimenional (3-D) formats. EVs from T cells infected with an HIV-1 provirus, which contains a dysfunctional mutant HIV Tat and lack of the Nef gene, can still stimulate KSHV infection. Although HIV-associated (HIV+) EVs lack viral proteins that are involved in cellular processes; they contain the HIV TAR RNA in excess of other HIV RNAs. We demonstrate that TAR RNA alone and TAR RNA-bearing EVs are able to enhance KSHV infectivity in oral epithelial cells, indicating the importance of the HIV TAR RNA in promoting KSHV infection. HIV+ EV-enhanced KSHV infection is blocked by the monoclonal antibody against EGFR. Our findings reveal that HIV+ saliva EVs is a risk factor for enhancement of KSHV infection and that inhibition of EGFR serves as a novel strategy for potentially controlling KSHV infection and transmission in the oral cavity.

## Results

### HIV-associated EVs enhance KSHV infectivity in oral epithelial cells

We treated iSLK-BAC16 cells with sodium butyrate and doxycyline to produce infectious KSHV virions, which contain a green fluorescent protein (GFP) cassette for monitoring success infection in terget cells [30]. To determine the multiplicity of infection (MOI) for each of KSHV stocks, we infected the immortalized oral epithelial line OKF6/TERT2 [31] using each KSHV preparation at various dilutions for 24 hr, followed by immunofluorescent staining of cells for KSHV-specific markers. We found that KSHV infected OKF6/TERT2 cells, leading to expression of the KSHV latency-associated nuclear antigen (LANA) and GFP and that 1:100 dilution of the KSHV stocks was estimaed to equal to MOI 0.1, which was used throuout all the experiments to ensure consistant results (S1 Fig).

To determine whether HIV+ EVs were able to affect KSHV infection in oral epithelial cells, we incubated OKF6/TERT2 cells with KSHV virions in the presence of EVs isolated from culture supernatants of latently HIV-1-infected J1.1 T cells or control “virus-free” Jurkat cells [26,32]. HIV+ EVs from J1.1 cells significantly enhanced KSHV infection compared to those isolated from control Jurkat T cells as determined by immunofluorescence microscopy of LANA and GFP proteins (Fig 1A) and flow cytometry on GFP of infected cells (Fig 1B and 1C), suggesting that HIV+ EVs potentially stimulated KSHV infectivity in oral epithelial cells. KSHV infects the oral cavity and oropharynx and the infection is more prevalent in HIV-positive people than that in the general population [6,33]. We postulated that saliva EVs in people living with HIV might be responsible for the higher KSHV infection in HIV patients. To test this hypothesis, EVs were purified from the saliva of HIV-infected donors and healthy individuals using the differential ultracentrifugation protocol [26]. To determine whether EVs prepared from saliva met the minimal requirement for EVs [34], we conducted immumoblotting on total saliva EVs. Saliva EVs from both HIV+ and HIV-donors contained tetraspanin proteins CD63, CD9 and CD81 (Fig 2A), suggesting presence of exosomes in the EV preparations [34]. In addition, the total EV protein amount was proportionally increased as more saliva was used in EV purifications (S2 table), indicating our protocol was able to purified EVs from saliva samples [34]. To determine whether HIV+ saliva EVs had HIV-specific components, we performed RT-PCR on total EV RNA and found that only HIV+ saliva EVs contained HIV TAR, Tat and Nef RNA, but not Env RNA (Fig 2B) [26], indicating that saliva of HIV-infected people contained HIV+ EVs. To evaluate the effect of saliva EVs on KSHV infection, we infected OKF6/TERT2 cells with KSHV in the presence of EVs from saliva of HIV-infected subjects and control people, respectively. Saliva EVs from HIV-infected subjects (Fig 2C, P8 and P9) significantly enhanced KSHV infection in OKF6/TERT2 cells compared to EVs from the saliva of healthy individuals (Fig 2C, N1 and N3) as determined by GFP flow cytometry. To determine whether KSHV was able to infect primary human oral epithelial cells (HOECs), we treated the cells with KSHV virions and found that KSHV-infected HOECs and infected cells expressed KSHV LANA and GFP (Fig 2D), indicating that KSHV can infect primary oral epithelial cells. To test whether HIV+ EVs were able to stimulate KSHV infection in these cells, we treated HOECs with EVs from J1.1 and Jurkat cells as well as those purified from saliva of HIV+ or HIV-donors. EVs isolated from HIV+ T cells or purified from the saliva of HIV-infected donors significantly stimulated KSHV infection in HOECs compared to EVs from control T cells and saliva of healthy donors, respectively (Fig 2E). To verify the stimulatory effect of HIV+ saliva EVs on KSHV infection, we treated HOECs with saliva EVs derived from HIV-infected donors (*n*=8), who were under ART treatment with CD4^+^ T-cell counts over 200 per ml, or those from healthy individuals, followed by the KSHV infection assays. HIV+ saliva EVs considerably enhanced KSHV infection compared to HIV-saliva EVs in primary HOECs (Fig 2F and 2G). Collectively, these results demonstrate that HIV+ saliva EVs indeed promote KSHV infection in oral epithelial cells.

**Fig 1.**
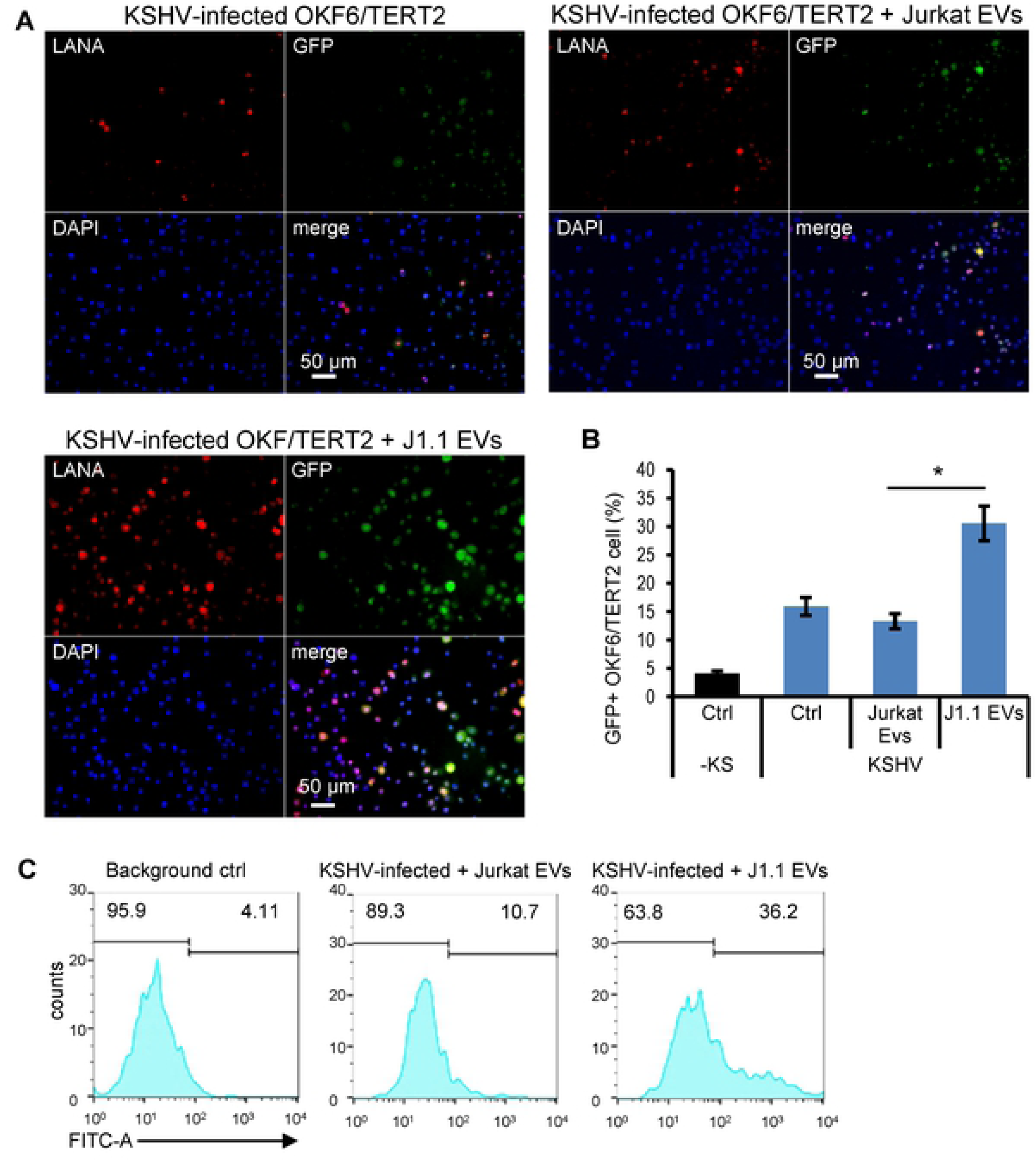
HIV+ EVs promote KSHV infection in oral epithelial cells. (A) OKF6/TERT2 cells were treated with EVs from J1.1 (HIV+) or Jurkat (HIV-) cells at 4 × 10^9^ EVs ml^-1^ [34], followed by KSHV infection for 20 hr. Cells were fixed for immunofluorescent staining using antibodies to LANA and GFP. (B) Flow cytometry of GFP+ cells after KSHV infection in the presence of EVs from J1.1 and Jurkat cells, respectively. Ctrl, KSHV alone. Data represent one independent experiment (*n*=4) out of three repeats. \**p*<0.05, *F*-test. (C) Flow cytometry histogram of (B).

**Fig 2.**
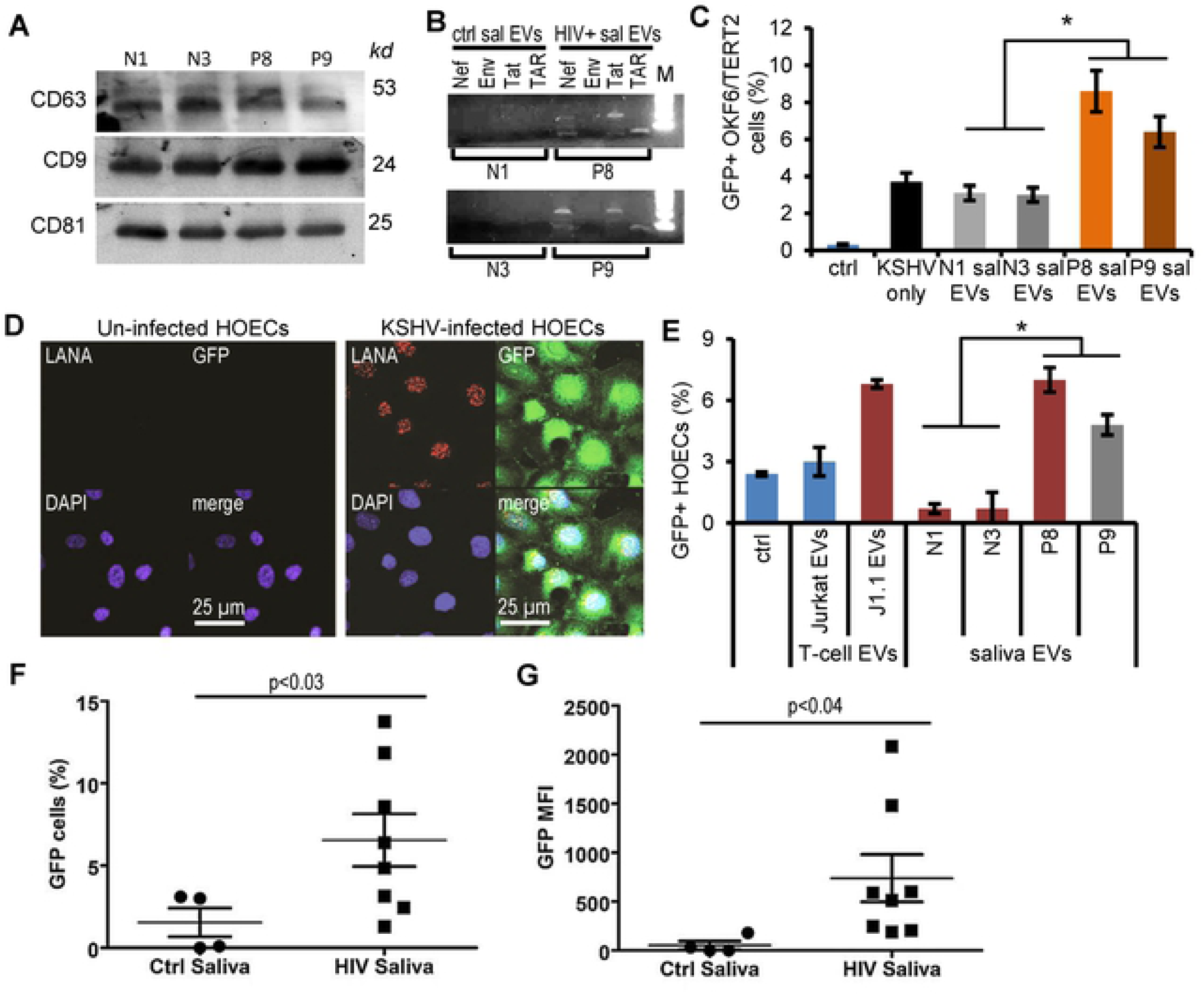
HIV+ EVs from saliva of HIV-infected donors promote KSHV infection in oral epithelial cells. (A) Immunoblots of total proteins extracted from saliva EVs from healthy (N1 and N3) and HIV+ (P8 and P9) individuals. Molecular weight for each protein indicated. (B) RT-PCR on total RNA extracted from saliva EVs of healthy and HIV+ donors. M, DNA size marker. (C) OKF6/TERT2 cells were treated with HIV+ (P8 and P9) and HIV-(N1 and N3) saliva EVs (100 μg ml^-1^), respectively, and then infected with KSHV for 20 hr. Infection was quantified by GFP flow cytometry. *, *p*<0.05. (D) LANA (red) and GFP (green) expression in primary human oral epithelial cells (HOECs) upon KSHV infection using immunofluorescent staining. nuclei, blue (DAPI); representative images shown. (E) HOECs were treated with saliva EVs (100 μg ml^-1^) from healthy (N1 and N3) and HIV+ (P8 and P9) donors, followed by KSHV infection. KSHV-infected GPF+ HOECs were quantified using flow cytometry. Data represent mean ± S.D. *, *p*<0.05. (F). HOECs were treated with saliva EVs (100 μg ml^-1^) purified from healthy (n=4) and HIV-infected donors (n=8), followed by KSHV infection. KSHV-infected GFP+ HOECs were determined by flow cytometry. \**p*<0.03, one-way ANOVA. (G) Mean fluorescence intensity plot of (F). \**p*<0.04, one-way ANOVA.

### HIV+ saliva EVs stimulate KSHV infection and transmission in 3-D culture models of oral mucosa

The oral mucosa is the first target of KSHV infection once the virus is in the oral cavity [6,7,8,9]. To evaluate the initial infection process of oral mucosa by KSHV, we created the 3-dimentional (3-D) organotypic culture by using OKF6/TERT2 cells as described previously [35] (Fig 3A), followed by KSHV infection. KSHV infected all layers of cells of the 3-D cultures of OKF6/TERT2 cells in the presence of HIV+ EVs from J1.1 T cells; however, the infection was barely detected in the presence of Jurkat cell EVs (Fig 3B). Further, we used 3-D cultured oral buccal mucosal tissues consisting of primary human oral epithelial cells (MatTek Co., Ashland, MA) (Fig. 3*A*, lower panel). EVs from HIV+ J1.1 T cells increased expression of KSHV LANA protein (Fig 3C, arrowheads) and GFP (Fig 3C, arrows) upon KSHV infection in the mucosal tissues and cells in the basal layers compared with HIV-EVs from Jurkat cells. Quantification of Fig 3C demonstrated that HIV+ J1.1 T-cell EVs significantly increased KSHV-infected LANA-expressing cells compared to HIV-Jurkat T-cells EVs (Fig 3D). Our findings suggested that HIV+ EVs were able to facilitate KSHV transmission through oral mucosa.

**Fig 3.**
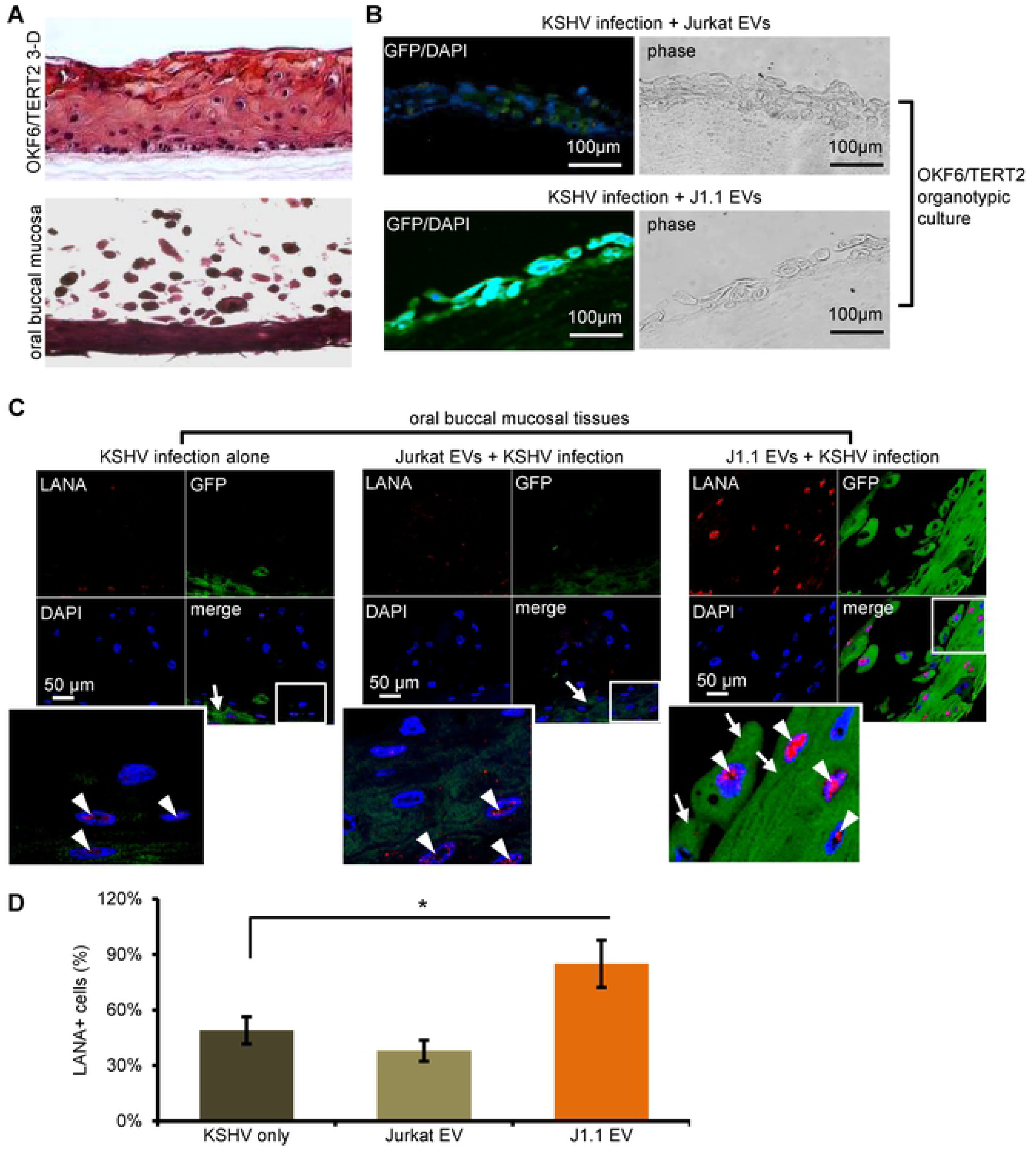
KSHV infection is increased by HIV+ EVs in 3-D cultural models of oral epithelial cells. (A) Haemotoxylin and eosin (H&E) staining of the organotypic culture of OKF6/TERT2 cells (upper panel) and the oral buccal mucosal tissue consisting of primary human oral epithelial cells (MetTak Inc.). Representative images shown. (B) HIV+ J1.1 T-cell EVs promote KSHV infectivity in a 3-D organotypic culture model constructed using OKF6/TERT cells. GFP+ cells represent KSHV-infected cells. Representative images shown. (C) The 3-D cultured oral buccal mucosal tissues were treated with EVs from HIV-infected J1.1 and control Jurkat T cells, respectively, followed by KSHV infection. Tissue sections were stained with antibodies to LANA and GFP. Red, LANA; green, GFP; blue, nuclei. Arrows, GFP; arrowheads, LANA. Representative images shown. (D) Quantification of LANA+ cells vs. total cells of (C). Data represented mean ± S.D.

### HIV transactivation element RNA (TAR) is critical for promoting KSHV infection

We suspected that HIV-specific EV cargo components were responsible for HIV+ EV-enhanced KSHV infection in oral epithelial cells. Latently HIV-infected J1.1 T-cell EVs do not contain HIV-1 viral particles, although these EVs have viral proteins such as Gag and the precursor form of Env protein (p160) [24]. To determine if HIV+ EVs contained viral proteins, such as Tat and Nef that are known to contribute to cellular functions [36,37], we performed immunoblot on total EV proteins isolated from latently HIV-infected J1.1 cells and the HIV+ Jurkat clone C22G cell line that contains a disruptive HIV *tat* mutant and *nef* deletion [38]. While the whole protein lysates of HIV-infected J1.1 cells contained Tat and Nef proteins, EVs from J1.1 and C22G cells did not produce the HIV proteins (Fig 4A), suggesting that the HIV-associated proteins might not play a major role in promoting KSHV infection. We have reported that EVs from both J1.1 and C22G cell lines contain HIV TAR RNA, which is required for HIV+ EV-enhanced proliferation of cancer cells [26]. HIV+ saliva EVs also contained TAR RNA (Fig 2B). Therefore, we postulated that the TAR RNA-bearing EVs contributed to increase KSHV infection in oral epithelial cells. To test this hypothesis, we treated OKF6/TERT2 cells with EVs derived from J1.1 and C22G cells, respectively, followed by KSHV infection. HIV+ EVs from both J1.1 and C22G cell lines promoted KSHV infectivity in OKF6/TERT2 cells as shown by GFP flow cytometry (Fig 4B), suggesting that the TAR RNA contributed to HIV+ EV-enhanced KSHV infectivity. HIV TAR RNA can directly induce expression of pro-oncogenes and proliferation of cancer cells and the pro-tumor effect of TAR RNA requires the bulge-loop structure [39] of the molecule [26]. To determine whether the bulge-loop structure of the TAR RNA affected KSHV infection, we transfected OKF6/TERT2 cells with a TAR RNA mutant containing 5 nucleotide replacements in bulge and loop sequences [26] and found that, while the wild type TAR RNA promoted KSHV infection in oral epithelial cells, the mutant TAR RNA failed to affect KSHV infection (Fig 4C, TAR vs. mut TAR). In addition, the RNA aptamer R06, which is complementary to the TAR apical region and blocks TAR function without disrupting the secondary structure of TAR [40], blocked TAR RNA-enhanced KSHV infection in OKF6/TERT2 cells (Fig 4C, TAR+R06). However, a scrambled aptamer [26] did not significantly change TAR RNA-induced KSHV infectivity (Fig 4C, TAR+scrb). These results indicate that the bulge-loop region of TAR RNA is critical for its function associated with the enhancement of KSHV infection in oral epithelial cells by TAR RNA-bearing EVs.

**Fig 4.**
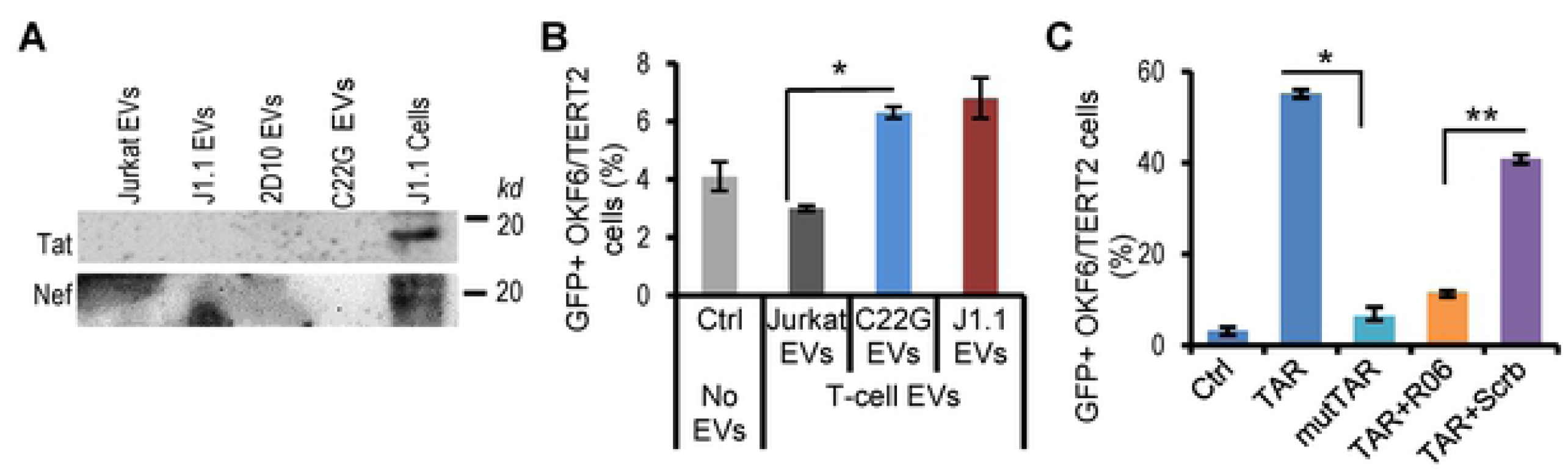
KSHV infection is enhanced by the HIV TAR RNA. (A) Immunoblots of HIV Tat and Nef proteins on total proteins extracted from EVs isolated from cultural supernatants of Jurkat, J1.1 and C22G cells. Total cell lysates of J1.1 cells (J1.1 Cells) were used as control. (B) Infection of KSHV in OKF6/TERT cells in the presence of EVs from Jurkat, C22G and J1.1 T cells. GFP+ KSHV-infected cells were determined by flow cytometry. *n*=3; *, *p*<0.05; *F*-test. (C) OKF6/TERT2 cells were transfected with synthetic HIV TAR RNA (TAR), the mutant TAR RNA (mutTAR), TAR RNA together with the R06 aptamer (TAR+R06) or the scrambled aptamer (TAR+Scrb), followed by KSHV infection for 20 hr. KSHV transfection was determined by GFP flow cytometry. Data represent one independent experiment (*n*=4) out of three repeats. \**p*<0.01, \*\**p*<0.02, *F*-test.

### HIV-associated EVs promote KSHV infectivity in an EGFR-dependent fashion

We have reported that EVs released from HIV-infected T cells and purified from plasma of HIV-positive patients stimulate proliferation of HNSCC and lung cancer cells in an EGFR-dependent manner through phosphorylation of ERK1/2 [26]. We reasoned that HIV+ EVs might promote KSHV infection in oral epithelial cells via the similar mechanism. Indeed, treatment of OKF6/TERT2 cells with cetuximab, a monoclonal antibody that blocks ligand binding to EGFR, inhibited KSHV infection in OKF6/TERT2 cells as shown by reduced numbers of GFP+ cells in the culture (S3 Fig, HIV+ J1.1 EVs vs. +cetuximab). The inhibitory effect of cetuximab on HIV+ EV-enhanced KSHV infection in OKF6/TERT2 cells was also determined by flow cytometry (Fig 5A). To determine whether viral proteins defining KSHV productive infection were affected by inhibition of EGFR, we treated OKF6/TERT2 cells with cetuximab and AG1478, a selective inhibitor of EGFR phosphorylation [41], followed by KSHV infection assays in the presence and absence of HIV+ EVs. HIV+ EVs enhanced expression of viral LANA and K8 proteins, an early viral protein encoded by open reading frame K8 to regulate viral and host cell transcription [42,43]. However, expression of the viral proteins was blocked by cetuximab and AG1478 (Fig 5B). Cetuximab also blocked HIV+ EV-enhanced KSHV infection in primary oral epithelial cells (Fig 5C). To determine whether inhibition of EGFR affected KSHV infection promoted by HIV+ saliva EVs in oral mucosal tissues, we treated the 3-D oral mucosal tissues with cetuximab, followed by KSHV infection in the presence of HIV+ saliva EVs. While HIV+ saliva EVs stimulated KSHV infection in oral mucosal tissues, cetuximab blocked the pro-infection effect of HIV+ saliva EVs in the tissues (Fig 5D). Our findings indicate that blocking EGFR was able to inhibit KSHV infection mediated by HIV+ EVs in the oral cavity.

**Fig 5.**
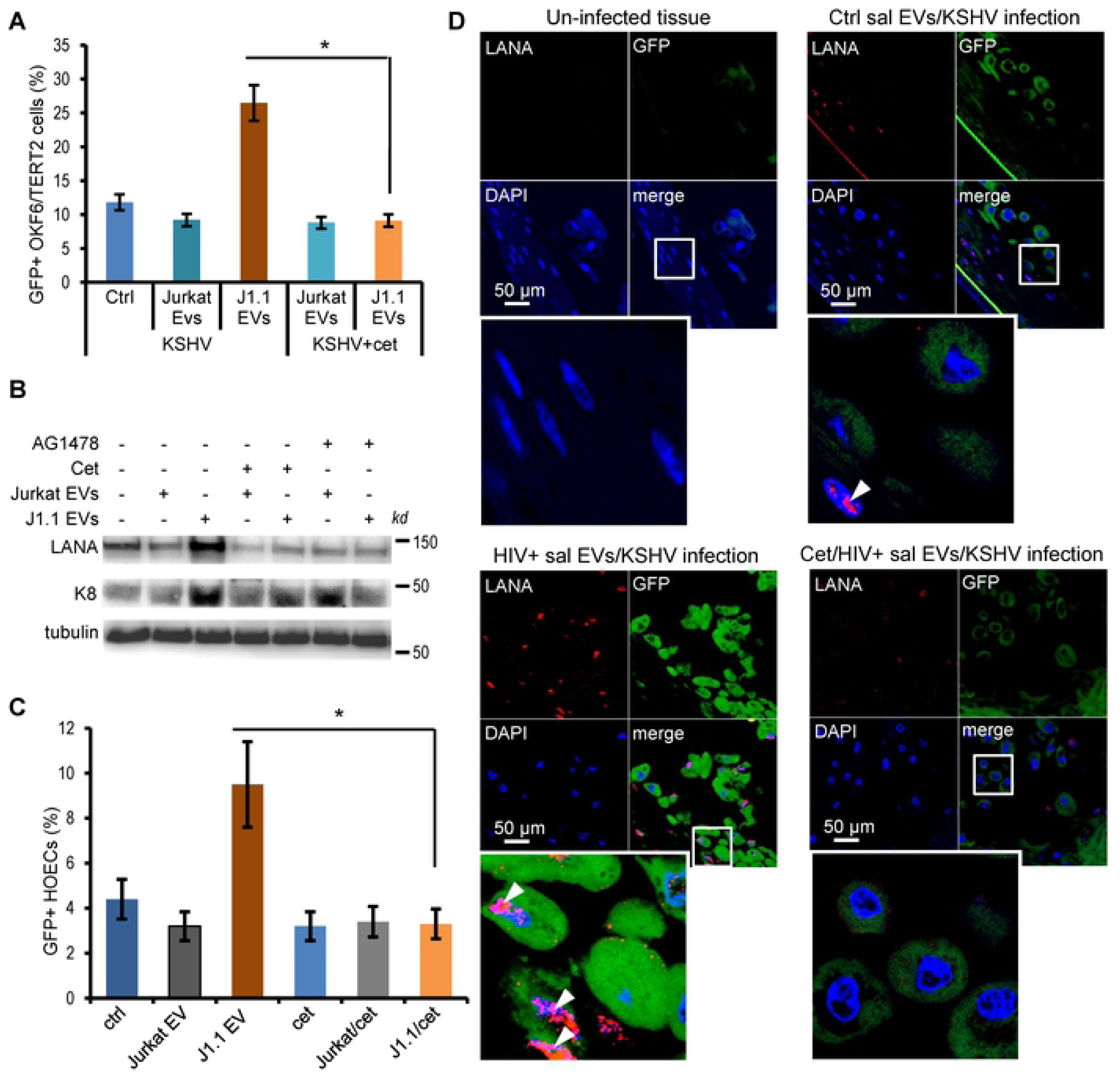
Increased KSHV infection by HIV+ EVs is EGFR dependent. (A) KSHV infected in OKF6/TERT2 cells in the presence of EVs from Jurkat or J1.1 cells (4 × 10^9^ EVs ml^-1^) with or without cetuximab treatment (20 μg ml^-1^). Data (mean±S.D.) represent one independent experiment (*n*=3) out of three repeats. \**p*<0.02, *F*-test. (B) OKF6/TERT2 cells were pre-treated with J1.1 and Jurkat EVs (4×10^9^ ml^-1^), respectively, in the presence or absence of cetuximab (Cet) or AG1478 (2 μm) for 30 min, followed by KSHV infection for 20 hr. Total protein lysates of cells were used for immunoblotting on viral LANA and K8 proteins. (C) Flow cytometry of GFP+ KSHV-infected OKF6/TERT2 cells treated with EVs from Jurkat and J1.1 cells treated with or without cetuximab (20 μg/ml). *n*=3; *, *p*<0.05; *F*-test. (D) Oral buccal mucosal tissues were treated with J1.1 or Jurkat cell EVs (4×10^9^ EVs ml^-1^) with or without cetuximab (cet), followed by KSHV infection. Arrowheads, LANA; green, GFP; blue, nuclei.

### HIV-associated EVs stimulate p38 MAPK signaling through EGFR

We have reported that HIV+ EVs induce phosphorylation of ERK1/2 in an EGFR-dependent manner without causing activation of the receptor in cancer cells [26]. To determine if HIV+ EVs contribute to activation of EGFR and its down-stream effector kinases, we treated OKF6/TERT2 cells and HOECs with EVs isolated from HIV+ J1.1 T cells and control Jurkat cells, respectively. Treatment of OKF6/TERT2 cells with HIV+ EVs for 10 min induced phosphorylation of p38 MAPK, a process was blocked by cetuximab and AG1478 (Fig 6A). Similarly, HIV+ EVs induced phosphorylation of p38 MAPK, but not ERK1/2, in HOECs (Fig 6B). However, HIV+ Evs failed to phosphorylate EGFR at tyrosine residuals 1068 (Y1068) and Y1173 in OKF6/TERT2 cells and HOECs, while EGF induced phosphorylation of EGFR at Y1068 and Y1173 as well as phosphorylation of p38 MAPK and ERK1/2 in OKF6/TERT cells and HOECs, indicating that the non-cancerous oral epithelial cells responded to EGF signaling (Fig 6A and 6B). In addition, HIV+ EVs and EGF failed to phosphorylate STAT3, a downstream effector in the EGFR signaling [44,45]. Our findings suggested that HIV+ EVs enhanced KSHV infection in an EGFR-dependent manner possibly through activation of the EGFR/p38 signaling in oral epithelial cells.

**Fig 6.**
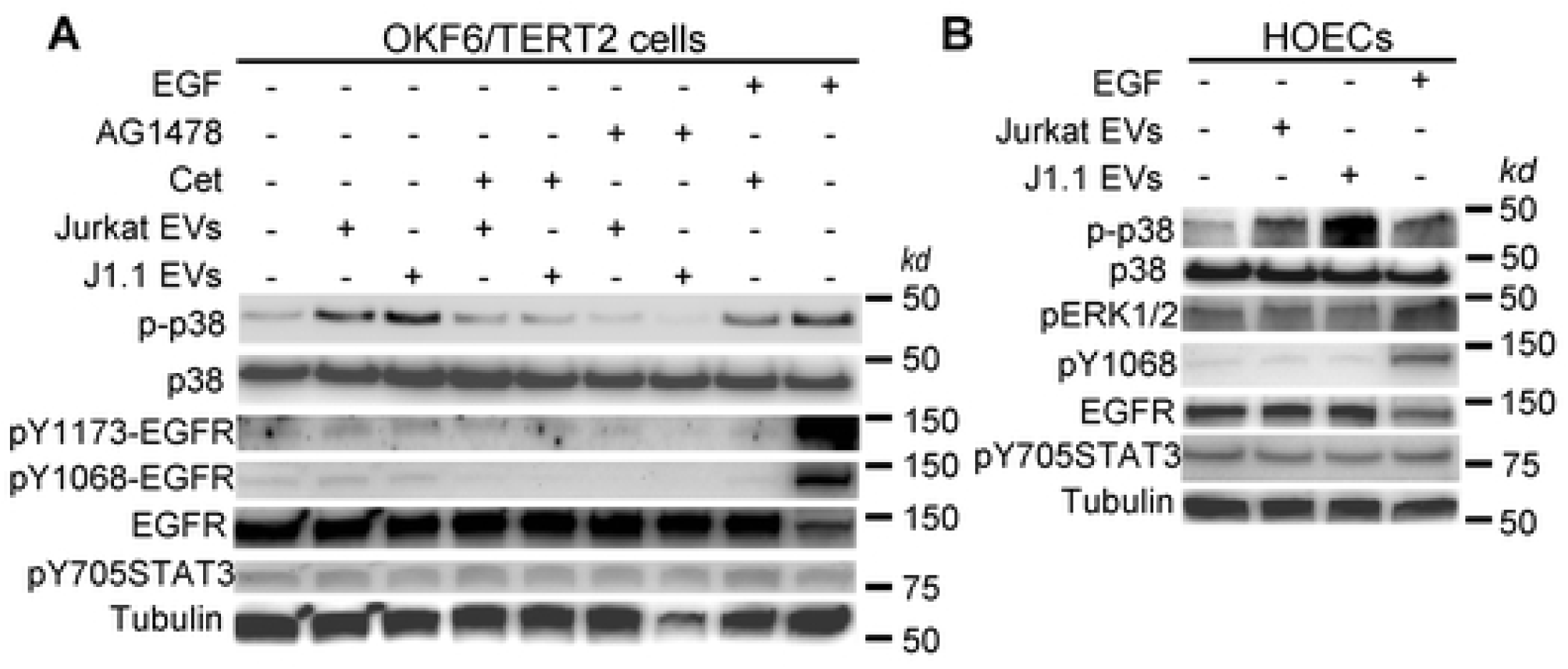
HIV+ EVs activate p38 MAPK via EGFR in oral epithelial cells. (A) OKF6/TERT2 cells were pre-treated with cetuximab (Cet, 20 μg ml^-1^) or AG1478 (2 μm) for 30 min, followed by treatment with J1.1 and Jurkat EVs (4 × 10^9^ EVs ml^-1^), respectively, for 10 min. Total protein lysates were used for immunoblotting. p-p38, phosphorylated p38; pY1173-and pY1068-EGFR, phosphoryled EGFR at 1173 and 1069 tyrosine residues, respectively. EGF (10 ng ml^-1^) was used as positive control. (B) HOECs were treated with HIV+ J1.1 or control Jurkat EVs (4x 10^9^ ml^-1^) for 10 min, followed by cellular lysis. Total cellular proteins were used for immunoblot. EGF (10 ng ml^-1^) treatment was used as a positive control.

## Discussion

HIV-infection is essential for KSHV co-infection, transmission, and its progression to malignancies [46]. In people living with HIV/AIDS, co-infection with KSHV is much more likely to lead to the development of KS and other KSHV-associated diseases [47,48,49]. The incidence rates of KSHV detection are more prevalent in the HIV-infected population than that in the general population in a case control study [33]. In this report, we demonstrate that HIV+ EVs from the saliva of HIV-positive patients and culture medium of HIV-infected T cells promote KSHV productive infection in oral epithelial cells cultured in both monolayer and 3-D models, indicating that HIV+ EVs are capable of regulating the initial steps of KSHV infection in the oral cavity. Saliva-mediated oral transmission of KSHV is considered as the most common route for spreading among homosexual people through deep kissing and “mother to child” transmission [3,4,5,6,7,8,9]. Because both saliva and peripheral blood samples [26] from HIV-infected persons contain HIV+ EVs, our findings suggest that the in HIV seropositive people bear higher risk for KSHV infection most likely through EVs in the body fluids.

It has been reported that oral microbial metabolites contribute to infection and lytic activation of KSHV [50,51,52]. Supernatants of periodontopathic bacteria cultures induce KSHV replication in BCBL-1, a latently infected KSHV-based lymphoma-derived cell line, and infection in embryonic kidney epithelial cells, human oral epithelial cells and umbilical vein endothelial cells [51,52]. The saliva of patients with severe periodontal disease contain high levels of short chain fatty acids that stimulate lytic gene expression of KSHV in a dose-dependent fashion in BCBL-1 cells [52]. These bacterial metabolic products can stimulate KSHV replication in infected cells using different mechanisms [51,52]. However, it is not clear whether these microbial metabolic products are responsible for KSHV infection in HIV-infected persons in the oral cavity. Collectively, our findings and these previous reports denote that multiple microbial/viral risk factor contribute to KSHV pathogenesis in the oral cavity.

We have reported that HIV+ EVs stimulate proliferation and proto-oncogene expression of squamous cell carcinoma cells in an EGFR-dependent manner [26]. Similarly, EGFR is critical for HIV+ EV-enhanced KSHV infectivity; blocking the receptor with a neutralizing antibody effectively inhibits KSHV infection in primary and immortalized oral epithelial cells. EGFR mediates HIV+ EV entry into target cells and participates in EV-induced signaling, including phosphorylation of ERK1/2, in head and neck as well as lung cancer cells [26]. However, cancer cells lacking EGFR, such as B-cell lymphoma cells, do not respond to HIV+ EVs [26]. Our results suggest that HIV+ EVs specifically promote KSHV infectivity through EGFR in epithelial cells in the oral cavity.

EVs from plasma of HIV-infected people and culture supernatants of HIV-infected T cells contain HIV TAR RNA in vast excess over all viral mRNAs [24,26]. In patients with virtually undetectable virion levels, TAR RNA can still be found in blood EVs [27]. Our results show that HIV+ saliva EVs contained TAR RNA and that synthetic TAR RNA considerably increases KSHV infection in oral epithelial cells. Several reports have shown that the HIV TAR RNA is a critical component of the HIV+ EV cargo and induces expression of proinflammatory cytokines and proto-oncogenes in primary human macrophages and head and neck cancer cells, respectively [24,26,27]. Synthetic TAR RNA alone can stimulate proliferation and migration of head and neck cancer cells [26]. The mutant TAR RNA with 5-nucleotide substitutions in the bulge and loop sequences fails to induce gene expression in head and neck cancer cells [26]. Similarly, our results demonstrate that the same TAR RNA mutant cannot enhance KSHV infection in oral epithelial cells. In addition, the R06 nucleotide aptamer, which creates an imperfect hairpin to complement to the entire TAR loop to block the function of TAR RNA [40], blocks TAR RNA-induced KSHV infection in oral epithelial cells. The R06 aptamer and its derivatives are able to reduce HIV-1 infection and inhibit the viral transcription [53,54]. Kolb *et al*. have reported that the replication of HIV-1 and the activity of β-galactosidase under the control of the HIV-1 5’LTR were reduced in cells expressing the nucleolar R06 transcript [53], suggesting the antiviral activity of the nucleotide aptamer. Our results implicate that the R06 RNA aptamer and its functional derivatives can be potentially developed as a strategy for controlling co-infection of the herpesvirus in the HIV-infected population.

We have reported that HIV+ EVs activate the ERK1/2 singling through the EGFR-TLR3 axis to induce proto-oncogene expression and proliferation of head neck and lung cancer cells [26]. However, our data show that HIV+ EVs specifically activate the MAPK p38, but not ERK1/2, through EGFR without inducing phosphorylation of the receptor in non-cancerous oral epithelial cells. Inhibition of the catalytic activity of the phosphorylated p38 blocks KSHV reactivation, possibly through reduction in a global H3 acetylation and phosphorylation [55]. Various chromatin-silencing mechanisms, including histone deacetylation, repressive histone methylation, and DNA methylation, lead to silence of the genomes of herpesviruses and HIV during latency [56]. Multiple short chain fatty acids, including butyric acid, propionic acid, isovaleric acid, and isobutyric acid, inhibit class-1/2 histone deacetylases (HDACs) for histone hyperacetylation, resulting in expression of genes associated with the fate of KSHV infection and viral reactivation [57,58,59]. The epigenetic modifications, particularly acetylation of histones, are required for maintenance of KSHV latency in classic and AIDS-associated KS tissues [59]. Taken together, our findings provide an insight into the mechanisms underlying HIV-specific components and co-infection of KSHV in people living with HIV/AIDS through the oral cavity. In addition, targeting the HIV TAR RNA and EGFR of oral epithelial cells may serve as novel approaches to control KSHV infection in the HIV-infected population.

## Materials and methods

### Ethical statement

For all human subject studies, written informed consent was obtained from all study participants according to protocol approved by the Human Subjects Institutional Review Board (IRB) at Case Western Reserve University and University Hospitals Cleveland Medical Center. Only de-identified human specimens were collected and used for this work.

### Cell cultures and 3-D organotypic cultures

The J1.1 cell line was obtained from the NIH AIDS Reagent Program. C22G cells were obtained from Dr. Karn (Case Western Reserve University). Jurkat cells were purchased from American Type Culture Collection (TIB-152, ATCC, Manassas, VA). These cells were maintained in RPMI1640 medium (HyClone Lab., Inc., Logan, UT) supplemented with 10% exosome-depleted FBS, which was prepared by ultracentrifugation of FBS (ThermoFisher Scientific, Waltham, MA) at 100,000 × *g* for 16 h at 4°C [26], followed by collecting supernatants without disturbing the pellet. Primary human oral epithelial cells (HOECs) were isolated from healthy patients who underwent third-molar extraction at School of Dental Medicine as previously described [60].

HOECs and immortalized OKF6/TERT2 human oral keratinocytes were maintained as previously described [31,61]. EpiOral™ oral mucosal tissues were purchased from MatTek Co. (Ashland, MA), which consist of normal human oral keratinocytes that are differentiated into tissues with a non-cornified, buccal phenotype. The 3-D organotypic cultures were constructed following previously published protocols by Dongari-Bagtzoglou and Kashleva [35]. Briefly, collagen gel cushion was prepared on ice from rat-tail type I collagen (Cat# Corning 354249, Thermo-Fisher) supplemented with 10% FBS in DMEM and antibiotics. Fibroblast gel layer was prepared by mixing 1 ml of NIH3T3 cells with the collagen gel as mentioned above. Culture inserts containing gel cushion and fibroblast gel layer were cultured for 4 day, followed by addition of OKF6/TERT6 cells to the center of the insert and cultured for 3 days. These inserts were then lifted and cultured in airlifting medium for 14 days with change of the medium every other day.

### Preparation of KSHV virions and EVs

EVs were prepared from cell supernatants by differential ultracentrifugation with filtration steps [26]. Briefly, cell culture media were centrifuged at 400 × *g* for 5 min to remove cells, followed by centrifugation at 11,000 × *g* for 10 min to remove any possible apoptotic bodies and large cell debris. EVs were precipitated at 100,000 × *g* for 90 min at 4 °C (50.2Ti rotor, Beckman Coulter, Brea, CA) and suspended in PBS. Isolated EVs were quantified using the acetylcholinesterase (AChE) assay system [26] (System Biosci. Inc/SBI, Palo Alta, CA) and maintained at −80 °C in DMEM for later use. To purify EVs from saliva, 2 ml of saliva was centrifuged at 400 × *g* for 15 min to remove cell contaminants. After centrifugation at 11,000× *g* for 10 min, saliva EVs were pelleted by ultracentrifugation at 100,000 × *g* for 90 min at 4 °C (Optima™ Max-XP, Beckman Coulter). The EVs were washed in 2 ml PBS and pelleted again at 100,000 × *g* for 90 min and suspended in PBS. Saliva EVs were quantified using BCA assays to measure total EV proteins following the manufacture’s protocol (SBI).

### Flow cytometry analysis

OKF6/TERT or HOECs were washed 3 times with PBS, then suspended in 100 μl of PBS. Flow cytometric analysis was performed by Green Fluorescent Protein (GFP) on FACSAria Flow Cytometer (BD Biosciences). FACS data was analyzed with FlowJo software (TreeStar Inc.).

### RT-PCR and immunoblot

Total RNA was isolated and purified using High Pure RNA Isolation Kit (Roche) according to the manufacturer’s instructions. Extracted RNA was reverse-transcribed to cDNA (High-Capacity cDNA Reverse Transcription, Applied Biosystems). Regular PCR analysis was performed using Q5 Hot Start High-Fidelity 2x Master Mix (New England Biolabs) and detected by the T100™ Thermal Cycler (Bio-Rad). Sequences of primers are listed in *SI Appendix* table S1. For immunoblotting, total EV proteins were purified using the Total Exosome RNA & Protein Isolation Kit (ThermoFisher) following the manufacturer’s instructions. To prepare total cellular proteins, cells were washed with PBS and then cellular lysates were obtained by adding 300 μl of RIPA Lysis and Extraction Buffer (ThermoFisher). Protein lysates were separated by SDS-PAGE and then transferred onto polyvinylidene fluoride membranes (PVDF, Merck Millipore) for immunoblot analysis. Antibodies used in immunoblotting are listed in S4 Table. Protein detection was performed by chemiluminescence using an ECL kit (ThermoFisher) with the ChemiDoc XRS+ Imaging System (Bio-Rad).

### Immunofluorescence microscopy

Immunofluorescence microscopy of 3-D cultures were performed as previously described [62] with minor modifications. Briefly, each section (5 μm) was de-paraffinized 3 times in Clear-Rite™ 3 and hydrated with 100% Alcohol followed by 95% Alcohol. Samples were blocked with 10% donkey serum at room temperature for 1 hr. Each section was incubated with the primary antibody at 4°C overnight. After washing in PBS, sections were stained with the appropriate AlexaFluor-conjugated secondary antibody to the species of the primary antibody. Sections were then mounted with the VECTASHIELD Fluorescent Mounting Media (Vector Lab Inc., Burlingame, CA) containing DAPI to visualize nuclei. Immunofluorescent images were generated using AMG EVOS FL digital inverted fluorescence microscope (AMG, Mill Creek, WA). Confocal images were acquired with a Leica TCS SP8 system (Leica Microsystems) using a 63×/1.4 objective at a pixel size of 90 nm. Channels were acquired sequentially by line. For immunocytochemistry, cells on 8 well culture slide were fixed with 100% methanol at −20°C for 20 minutes followed by permeabilization with 0.3% Triton X-100 in PBS. Cells were then stained with the primary antibody followed by incubation with appropriate secondary antibodies. Fluorescent images were taken as described above. Antibodies for immunofluorescence microscopy are listed in S5 Table.

### Statistics

Results of treatments were compared with those of respective controls. Data are represented as mean ± S.D. Flow cytometry data were subjected to one-way ANOVA when sample sizes were *n ≤* 3. Statistical significance was considered at *p*< 0.05. For dada with *n ≤* =5, F-test was applied. Data analyses were performed and graphs were generated using Prism (GraphPad Software, La Jolla, CA) and Excel 2013 (Microsoft).

## Acknowledgments

We thank the Light Microscopy Image Core at Case Western Reserve University School of Medicine, supported by the NIH Office of Research Infrastructure Program Grant (S10-OD024996), and the Cytometry & Imaging Microscope Shared Resources at the Case Comprehensive Cancer Center, supported in part by the NIH/National Cancer Institute (NCI) grant (P30CA043703), for technical assists. The following reagents were obtained through the National Institute of Health (NIH) AIDS Reagent Program, Division of AIDS, National Institute of Allergy and Infectious Diseases (NIAID), NIH: HIV-1 LAV infected Jurkat E6 cells (J1.1) from Dr. Thomas Folks.

## Supporting information

**S1 Fig. Titration of KSHV infection in immortalized OKF6/TERT2 cells.** OKF6/TERT2 cells were grown in 6-well plates to 70-80% confluency and then were added with KSHV virions at different dilutions. After 20 hr incubations, cells were washed with PBS and fixed in methanol at 4 °C for 30 min. Immunofluorescence microscopy was performed using antibodies to KSHV LANA and GFP. Red, LANA; green, GFP; blue, nuclei.

**S2 Fig. Fig. S2. Inhibition of KSHV infection enhanced by HIV+ EVs by cetuximab.** OKF6/TERT2 cells were treated with EVs from Jurkat and HIV+ J1.1 T cells (4 × 10^9^ EVs ml^-1^), or remained un-treated, in the presence or absence of cetuximab (20 μg ml^-1^), followed by addition of KSHV virions. Microphotographs of GFP+ cells were taken 20 hr after KSHV infection.

**S3 Table. Total proteins from EV stocks prepared from various volumes of the saliva**

**S4 Table. Primers used in this report**

**S5 Table. Antibodies used in this report**

